# The evolution of sequence specificity in a DNA binding protein family

**DOI:** 10.1101/2024.06.24.600358

**Authors:** Meghna Nandy, Madhumitha Krishnaswamy, Mohak Sharda, Aswin Sai Narain Seshasayee

## Abstract

Transcriptional regulation enables bacteria to adjust to its environment. This is driven by transcription factors (TFs), which display DNA site recognition specificity with some flexibility built in. TFs, however, are not considered essential to a minimal cellular life. How do they evolve? It has been hypothesized that TFs evolve by gaining specificity (and other functions) on a background of non-specific chromosome structuring proteins. We used the IHF/HU family of DNA binding proteins, in which IHF binds DNA in a sequence-specific manner, whereas HU binds more indiscriminately, to test this hypothesis. We show that HUβ has been present from the bacterial root, while both IHF subunits emerged much later and diversified in *Proteobacteria*, with HUα having possibly arisen from transfer events in *Gammaproteobacteria*. By reconstructing ancestral sequences *in-silico* on a rooted phylogeny of IHF/HU we show that the common ancestor of this family was probably HU-like and therefore non-specific in binding DNA. IHF evolved from a branch of HU after HU had substantially diverged. Various residues characteristic of IHFα and shown to be involved in specific sequence recognition (at least in *E. coli*) have likely been co-opted from preexisting residues in HU, while those residues of IHFβ have likely evolved independently, suggesting that each of the IHF subunits has undergone different trajectories to acquire their DNA binding properties.

## Introduction

Transcriptional regulation enables an organism to react to changing circumstances. In bacteria, this is primarily driven by transcription factors (TFs), which bind to sites on the DNA, usually upstream or just downstream of transcription start sites, and activate or repress the expression of proximal genes (Balleza et al., 2009; Browning & Busby, 2004, 2016). The activity of TFs is thus mediated primarily by their ability to recognize and bind to particular nucleotide motifs in DNA, a highly evolvable property (Shultzaberger et al., 2012). Unlike restriction-modification systems that bind to short sequence motifs with “exquisite” specificity (Saravanan et al., 2008; Vasu et al., 2012), TFs binding to their target sites allows for some flexibility in their DNA site recognition.

Though regulation of transcription by TFs is important to adaptation, it is not absolutely required for cellular life. For example, surveys of essential and conserved genes across genomes suggest that TFs are rarely conserved across a large phylogenetic breadth (Gil et al., 2004); for instance, bacterial endosymbionts rarely code for TFs. In bacteria, the need for TFs emerges and expands super-linearly with an increase in complexity as defined by the number of gene functions required by an organism (Cases et al., 2003; Molina & van Nimwegen, 2009). On the other hand, the very ability of a protein to bind to DNA non-specifically, for example, to compress it so that it could fit within the dimensions of a cell, is necessary for even the most basal cellular life. Thus, the question arises - where and when did TFs evolve?

Sandhya Visweswariah and Stephen Busby point out that global TFs, in addition to binding at target sites that mediate regulation of gene expression of proximal genes, often bind to a large number of low-affinity sites and bend DNA, not unlike nucleoid-associated proteins (Visweswariah & Busby, 2015). In fact, the boundary between a global TF and a nucleoid-associated protein (NAP) is blurry (Dorman et al., 2020). They used this observation to hypothesize that specific DNA site recognition in TFs may have evolved on a background of nonspecific DNA binding observed in NAPs. This idea, however, remains to be tested systematically against data. In a previous study, we provided evidence that non-TF relatives of TFs in some protein superfamilies appeared, on average, earlier than TFs on a eukaryotic phylogenetic tree (Dubey et al., 2025). However, this study did not directly test the evolutionary trajectory in terms of sequence diversification of a single protein sequence family, which is strictly required to test Visweswaraiah and Busby’s hypothesis. Such testing would require reliable sequence alignments containing both specific and non-specific DNA binding proteins, which can then permit ancestral sequence reconstructions.

Integration host factor (IHF) and Histone-like DNA-binding protein (HU) provide an interesting model system to evaluate this idea (Boubrik et al., 1991; Drlica & Rouviere-Yaniv, 1987; Friedman, 1988). Both proteins, at least in *E. coli* and several other bacteria, comprise two subunits α and β and bind to DNA as dimers. All the IHF/HU subunits are characterised by the presence of two *α* helices followed by five *β* strands forming a sheet and ending at an *α* helix. HU in many organisms, however, exists as a single subunit protein. Even in *E. coli*, both homo- and hetero-dimers of HU form, each with its own properties (Claret & Rouviere-Yaniv, 1997; Hammel et al., 2016). HU binds to DNA but has no known recognition motif and thus has little sequence specificity (Swinger & Rice, 2007), though in certain conformations, it can recognize kinked DNA structures and act like a TF at a few sites (Verma et al., 2023). On the other hand, IHF forms heterodimers in *E. coli* and binds DNA with high affinity to a consensus sequence motif (Craig & Nash, 1984) and operates as a global TF. Despite the variation in their DNA binding specificity, both proteins belong to the same, tightly alignable sequence family. It is believed that IHF evolved from HU (Visweswariah & Busby, 2015), but this belief is grounded in learned intuition rather than systematic evidence.

In an important prior study, Dey et al. performed an analysis of a tree of the IHF/HU family towards identifying key structural features distinguishing IHF and HU, such as hydrophobicity and salt bridge formation (Dey et al., 2017). In a book chapter, Munoz et al. also used a HU/IHF tree to identify residues associated with major clade diversifications (Muñoz et al., 2010). However, these studies do not systematically assess the family’s evolutionary trajectory. In our study, under the assumption that HU and IHF share the same common ancestor, we investigate the evolutionary trajectory of this sequence family, in particular, the evolution of DNA recognition involved in TF activity.

## Methodology

### Dataset curation

The primary dataset was a subset from a list of 10,575 prokaryotic genomes and proteomes annotated in the Web of Life (WoL) (Zhu et al., 2019). Only complete and scaffold genomes, which were either reference or representative genomes, were taken, and strain level redundancy was removed. This resulted in a final dataset of 6,116 prokaryotic genomes, which was used for subsequent analysis. These 6,116 genomes were further curated to the genus level (only one representative per genus) for the construction of a 16s rRNA species tree.

Sequences containing the IHF/HU domain (Pfam: PF00216) were picked up from the dataset using hmmsearch (Eddy, 2011) to give a total of 12,184 proteins, which were classified on the basis of their KO annotation to each of the different subunits (K05787: HUα, K03530: HUβ, K04764: IHFα, K05788: IHFβ). This produced a total of 9,913 subunit sequences, which were trimmed to the length of the domain. The sequences were aligned using mafft (auto) (Katoh & Standley, 2013) and trimmed on the basis of excessive gap-containing columns. The sequences contributing to >95% of gaps in more than 5 residue positions were pruned from the alignment, and these sequences were not clade specific. A final list of 9,880 subunit containing domains were used to get a protein alignment of 9,880 subunit domains. This alignment was then used to run a PCA using sklearn in Python, using the algorithm described in (Konishi et al., 2019), to identify principal components (residues) contributing to the most variation in the sequences.

To supplement the alignment based analysis and classification, a structure based alignment using MAFFT-DASH (Rozewicki et al., 2019) was performed for a subset (N=990) of the 9,880 IHF/ HU sequences identified, and the two subset alignments compared using a PCA and NJ.

#### Species tree

The dataset of 6,116 taxa was subset at the genus level to get 1,454 genera, each represented by a randomly picked extant species belonging to a particular genus, selected at random. The 16S rRNA sequences of these representative taxa was picked up from NCBI to make the primary species tree. The sequences were aligned using mafft (auto) and the tree was constructed using IQ-TREE (Nguyen et al., 2015), and rooted between bacteria and archaea (midpoint rooting).

To supplement the 16s rRNA species tree, a bacterial representative GTDB tree (Parks et al., 2020) was obtained by pruning the “bac120” marker gene species tree to taxa in our dataset, resulting in a species tree with 3100 taxa. Tree topology and Robinson Fould tests were done by pruning both of the trees to list of common taxa (953 taxa), and tests were carried out using *ape* (Paradis & Schliep, 2019) and *phangorn* (Schliep, 2011) packages from R.

#### Ancestral State Reconstruction

The 16s rRNA species tree was used for subsequent maximum-likelihood ancestral state reconstruction analysis based on the presence/absence of different subunits in extant genus tips. The reconstruction was performed using the ace (discrete traits with marginal = TRUE) function from the *ape* package in R. Ancestral state estimation were then visualized in iTOL (Letunic & Bork, 2024) as piecharts for each internal node. The ancestral state reconstruction was done in parallel for the GTDB tree with the same parameters.

#### Gene tree

To construct a gene tree, the protein alignment was first converted to a codon alignment using PAL2NAL (Suyama et al., 2006). ModelFinder (Kalyaanamoorthy et al., 2017) was run with a sampled alignment to identify the best model (KOSI07) to be used for the entire gene tree (Kosiol et al., 2007). An unrooted gene tree was constructed with 1000 ultrafast bootstrap (Hoang et al., 2018) parameters. Testing of different outgroups was done on sampled alignments to reduce the computational load, by specifying outgroups during tree construction.

Phylogenetic reconciliation of the 16S genus level species tree with the corresponding unrooted gene tree was done using the OptRoot program in Ranger-DTL 2.0 (Bansal et al., 2018), with different variable cost parameters. OptRoot employs the method of phylogenetic reconciliation to identify a possible rooting of the gene tree that minimizes the costs of duplication, transfer and speciation. The threshold transfer costs, which are distance dependent, can also be varied across a range. OptRoot was run 100 times over a range of transfer costs using SummarizeOptRootings.

TraM Relaxosome protein from *Escherichia coli* (UniProt: P10026) was selected to be the outgroup due to consistent rootings with the sampled alignment, and hence, was added to the 9,880 genes and realigned to use in constructing the gene tree. The gene tree with the outgroup was constructed using IQ-TREE 2 (Minh et al., 2020).

Phylogenetic reconciliation of the 16S rRNA tree and the appropriately pruned gene tree was done using Ranger-DTL to estimate the number of duplications, transfers and speciation events. 100 reconciliations performed with default parameters were combined using AggregateRanger to give a final mapping of nodes experiencing duplication, transfer and speciation events.

#### Ancestral sequence reconstruction

The parameters for construction of the gene tree using IQ-TREE 2 specified an outgroup (-out) and the reconstruction of ancestral sequences (-asr). The “state” file output from IQ-TREE 2 runs gives the possible ancestral states at each codon position for each ancestral internal node, and this was then combined to give the ancestral sequences for each node. The codon sequences were translated to give protein sequences and key nodes identified from the gene tree (ancestral to a subunit class/overall ancestor) were appended to the protein alignment and used for further analysis.

A Neighbour Joining (NJ) tree was constructed using FastTree (Price et al., 2010) (using -noml) for characterization of the key ancestral nodes.

#### Shannon’s Entropy

Conserved sites were identified by taking subsets of each subunit class from the multiple sequence alignment. Shannon’s entropy was calculated at each column of the alignment with <50% gaps. Sites which had an entropy below the 1^st^ quartile of the overall distribution were taken to be conserved sites. The major amino acid occupying that site was obtained from viewing the alignment in Jalview (Waterhouse et al., 2009). This was done separately for each class of subunits. Residues found to be relevant for DNA binding were picked up from literature for both IHF and HU subunits.

#### PSSM – Position specific scoring matrix

To get an estimate of the extent of coverage of key residues, a PSSM was computed for the reference “motif”. This was done by first making a position count matrix based on the residues in the motif and creating a position frequency matrix. Since the overall alignment showed low conservation and noisy frequencies of the residue positions, a pseudo count (0.0476) was computed on the basis of the background expected frequencies (0.05 for each amino acid) and used to populate the position frequency matrix, with a gap penalty of -0.12. A log odds ratio of the position frequencies against the background frequencies was computed using the formula:

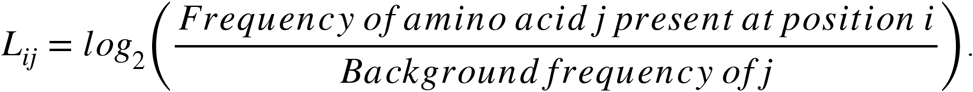

The PSSM for a “motif” (with n number of residues) is hence represented as a n *x* 21 matrix *L* (Log odds matrix), where the rows correspond to each position in the motif, and columns correspond to the 20 possible amino acids + a gap position. Each element of the matrix is a log odds score of the amino acid present at that particular position, represented by *L*_ij_ Given an alignment of length n, the PSSM score for a sequence *A* in the alignment is calculated as 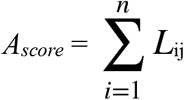 where *j* represents the amino acid present in *A* at residue position *i*.

The alignment files were trimmed to only constitute residues of interest using trimal (Capella-Gutiérrez et al., 2009), and scores were computed for each trimmed sequence. This scoring was done for both the extant as well as ancestor sequences on the basis of key residues for DNA binding in each subunit, and the key residues for specific base contacts in IHFα.

## Results

HU subunits are more widespread and present from the bacterial ancestor, while IHF subunits show late emergence

We searched a set of 6,116 bacterial and archaeal genome sequences for IHFα. IHFβ and HUα and HUβ, finally producing a dataset of 9,880 proteins (N = 6,648 for HUβ, 1,559 for IHFα, 1,537 for IHFβ and 136 for HUα). We performed a Principal Component Analysis (PCA) of the aligned sequences (Konishi et al., 2019), which showed that (a) HUα and HUβ cluster together without any clear separation between them; (b) IHFα and IHFβ are distinct from each other and from HU, consistent with previous work (Dey et al., 2017; Kamashev et al., 2017). The superset of HUα and HUβ subunit sequences is referred to as ‘HU’ in the downstream analyses. We also complemented the clustering with a Neighbour-Joining (NJ) tree, constructed from the complete alignment, which is consistent with the PCA map (Figure 1). Additionally, we also performed a structure-based alignment on a sampled set of ∼1,000 sequences. Both PCA and NJ-based clustering of sequences based on this structure-based alignment produced similar results (Supplementary figure S1).

**Figure 1.**
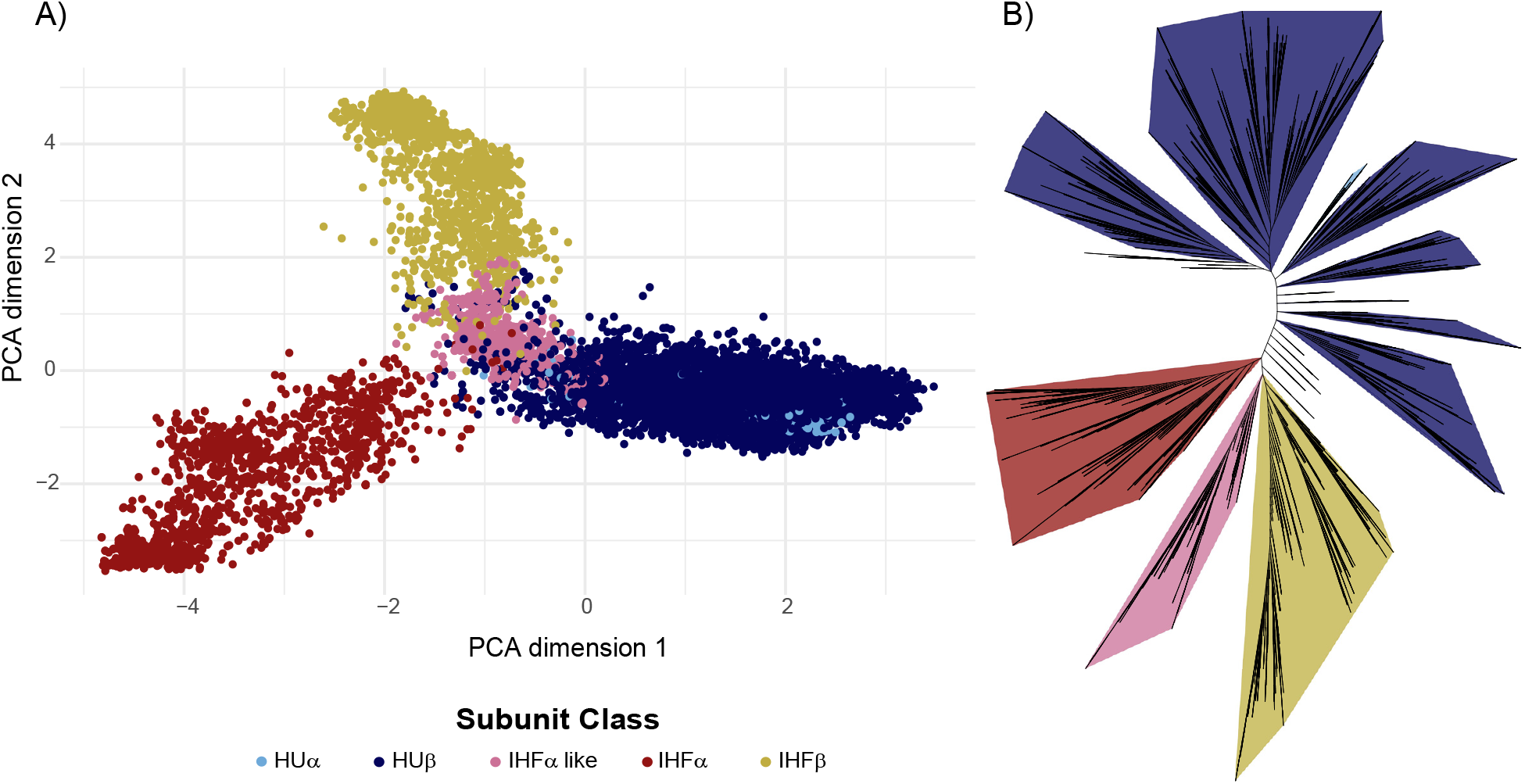
The IHF/HU sequence family. The clustering of the 9,880 IHF HU subunit sequences visualized using (A) Principal Component Analysis (PCA) based clustering.. Light blue circles: HUα, Dark blue circles: HUβ, Pink circles: IHFα like, Red circles: IHFα, Yellow circles: IHFβ. (B) Unrooted Neighbour Joining (NJ) tree of the sequences, constructed using FastTree and visualized, annotated using iTOL(Letunic & Bork, 2024)

A small subset of sequences annotated as HU in KO intermixed with IHF subunit proteins existing on the boundary between HU and IHFα on the PCA map. We analysed this subset of sequences more closely as follows. HU and IHF show distinct molecular weights and hydrophobicity. The subset of sequences of interest appears IHF-like on molecular weight but HU-like on hydrophobicity; however, on the gene tree (see below), this set clusters with IHFα. Based on these, we term this subset of proteins IHFα-like. Supplementary figure S2).

All the subunits are absent in most Archaea and are found predominantly in Bacteria. Within bacteria, 86% code for at least one of the four subunits. HUβ is the most conserved of the 4 subunits and is found in 81% of Bacteria (n = 4,695) with no clade-specific absence, while HUα is rarer and found in only 2% of organisms, predominantly *Gammaproteobacteria* (n=136). Both

IHF subunits are usually found together and in about 26% of the organisms (n = 1,538), mainly in *Proteobacteria*.

To understand how this conservation ties back to the emergence of the subunits, we needed a species tree to evaluate the gains and losses of all 4 subunits. Our primary representative species tree is a 16S rRNA tree representing 1,454 genera (see Methods). A more comprehensive representation of most of the 6,116 species (all bacteria) is a pruned, rooted GTDB (Genome Taxonomy Database) tree, representing 3,100 species from our dataset. The two trees show reasonable congruence (Mantel test: r = 0.4091, p = 0.001 and normalised Robinson Fould value: 0. 5516). We evaluated the emergence of the different subunits, based on their presence and absence in extant sequences, by performing an ancestral state reconstruction of the curated species tree (Fig. 2 and Supplementary Figure S3). HUβ appears to have emerged at the bacterial common ancestor and is present in almost all bacteria, with sporadic losses not specific to any clade. HUα is present a l m o s t e x c l u s i v e l y i n *Gammaproteobacteria*, appearing in a Gammaproteobacterial ancestor. IHFβ appears in the common ancestor of *Proteobacteria* and *Acidobacteria*, with IHFα appearing shortly after in the Proteobacterial lineage. IHFα-like emerges independently in a *Bacteroiidetes* ancestral node. These apply to both the 16S rRNA and the GTDB trees (Supplementary Figure S4).

**Figure 2.**
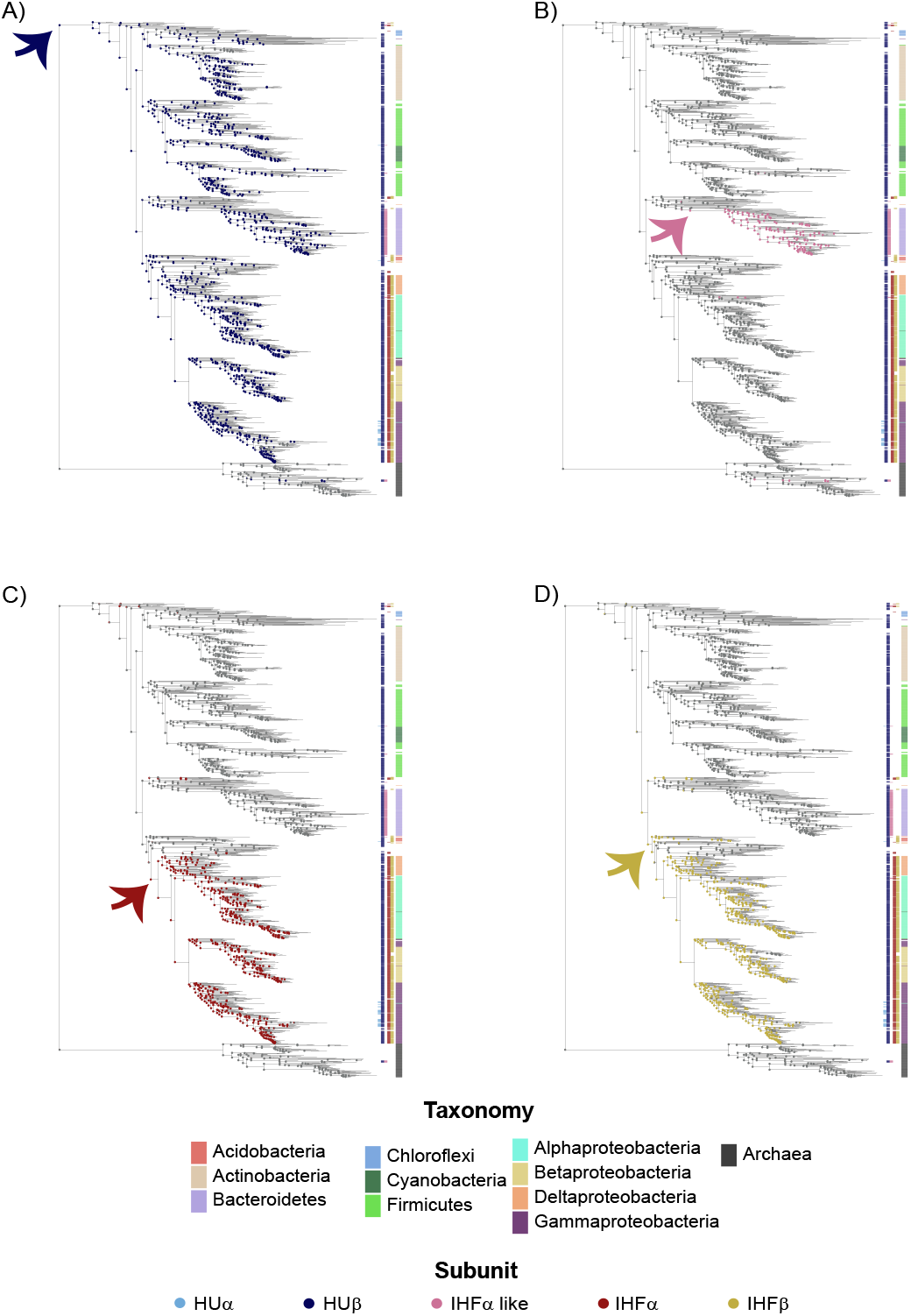
The bacterial species tree. 16S rRNA tree (with 1,454 tips) of genus level curation of taxa used in the dataset. The presence of the different categories of subunits in the ancestral nodes is shown as a pie chart for (A) HUβ (dark blue), (B) IHFα like (pink), (C) IHFα (red), (D) IHFβ (yellow). Coloured circles correspond to the possible presence of the different subunits in ancestral nodes while grey indicates absence or minimal likelihood of presence of the subunits. The curved arrows indicate node of first major emergence for each subunit, and colourstrips indicate presence/absence of different subunits along with their broad taxonomic annotation. The taxonomy of each of the species is colour coded to represent the major phyla or class. All trees visualized and annotated using iTOL.

Thus, HUβ has been present from the bacterial root, while both IHF subunits emerged much later and expanded in *Proteobacteria* and less so in other lineages.

### IHF/HU family ancestor resembles HU subunits

We explored several options for identifying a suitable rooting for the IHF/HU gene tree. We first constructed an unrooted tree and later rooted it at the midpoint. This placed a root between the HU subunits such that one clade consisted entirely of HU subunits, while the other consisted again of HU subunits, which later branched to give the IHF subunit clades. Upon subsampling of the gene tree, we noticed inconsistent results with midpoint rooting as it would place the root interchangeably between HU subunits and between different IHF subunits; hence, this method was unreliable. We then utilized phylogenetic reconciliation, which looks at identifying an optimal root by comparison of the species and gene trees and minimising the cost of duplication, transfer and loss events. This analysis also gave us inconsistent results with variable cost parameters as no single optimal root could be identified, and the set of all roots obtained included both IHF and HU subunits. (Supplementary Figure S5).

We finally asked if an outgroup-based rooting could provide us with consistent results that could be used for reliable evolutionary analysis. An initial rooting using other IHF/HU family members not annotated as IHF or HU did not work, as these clustered with the subunits (Supplementary Figure S6). We then examined the larger superfamily to which the IHF/HU family belongs to identify potential outgroups. This operates assuming that different families within a superfamily share a common ancestor. IHF and HU belong to a superfamily that comprises only one other family, the DNA binding domain of TraM, which is an essential component of the conjugative DNA transfer machinery; its structure is similar to that of IHF/HU (Supplementary Figure S7A). TraM clusters distinctly and away from the IHF/HU family and emerges as the only outgroup identifiable for HU and IHF (Figure 3A). This clustering holds true in the structure based alignment as well, with TraM separating out from the overall IHF/HU family (Supplementary Figure S7B). While we recognize the limitations of basing a phylogeny on a single outgroup, this appears to be a hard limit imposed by the sequence variation contained within this superfamily.

**Figure 3.**
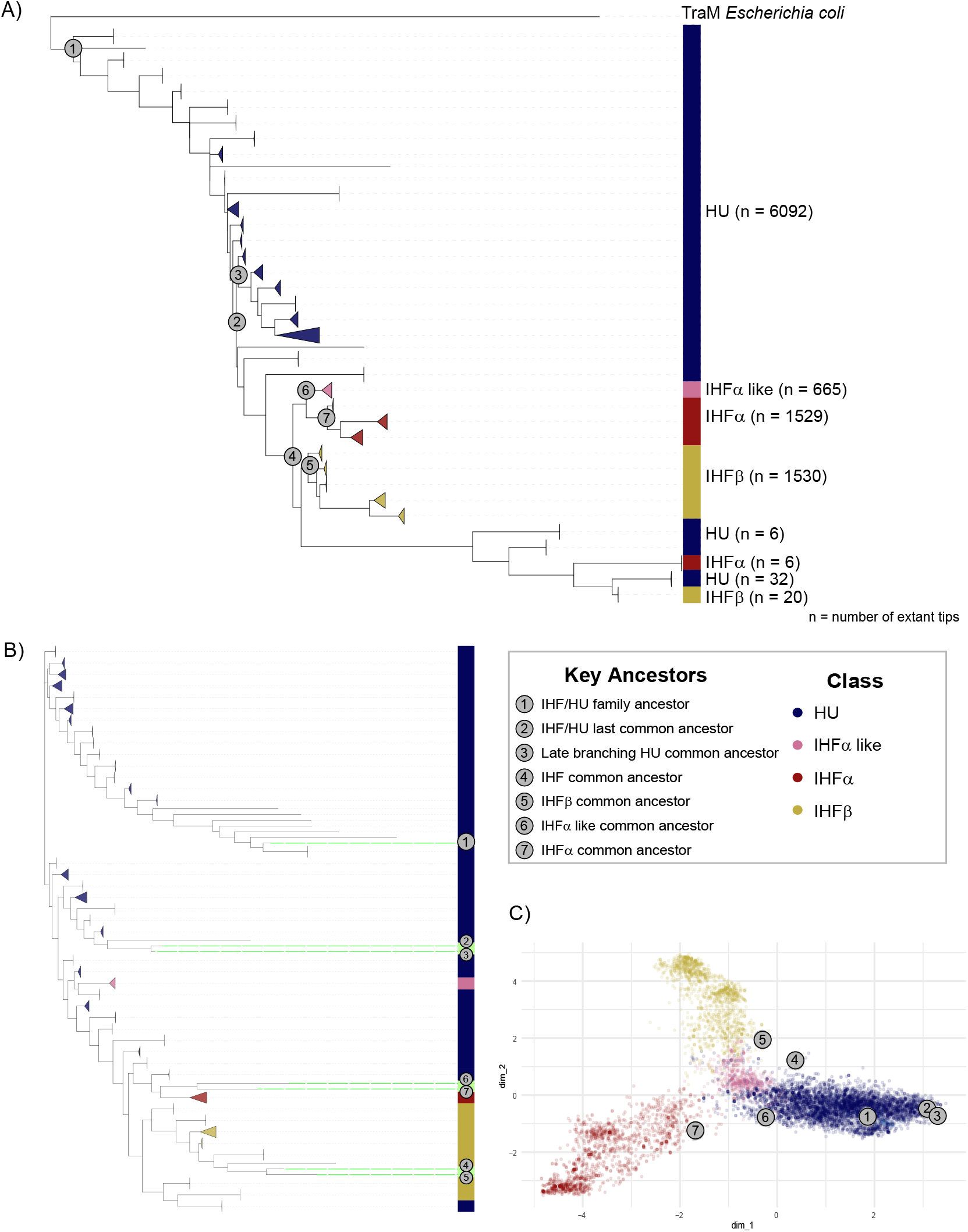
Gene trees and ancestral nodes. (A) A collapsed gene tree (with 9881 tips) showing TraM from Escherichia coli as the outgroup (visualized using iTOL), n values stand for number of sequences in the collapsed nodes, and size of the triangles indicate number of tips collapsed. (B) A collapsed Neighbour Joining (NJ) tree of extant sequences along with key ancestral nodes (C) PCA of extant sequences along with key ancestral nodes. Dark blue: HU (refers to the superset of both HUβ and HUα), Pink: IHFα like, Red: IHFα, Yellow: IHFβ, Grey circles with numeric annotations: key ancestral nodes.

We observed that the rooted gene tree shows a clear distinction of HU, IHFα and IHFβ, similar to the results of the PCA, with all the subunits segregating into clades with minimal intermixing. IHFα-like forms a monophyletic sister clade to IHFα. HUα sequences form two clades within the larger HUβ clade; however, HUα sequences, though found almost exclusively in *Gammaproteobacteria*, are not necessarily most closely related to Gammaproteobacterial HUβ sequences, suggesting a role for horizontal gene transfer in its formation (Supplementary Figure S8).

A protein related to HUβ appears to be the ancestor from which IHF subunits (α and β) have diverged. IHF and HU are not separate monophyletic clades; instead, IHF diverged after HU had already diversified to a great extent. To classify the ancestral nodes of the gene tree, we performed an ancestral sequence reconstruction of all the nodes of the gene tree and compared these to extant sequences using PCA and Neighbour joining-based clustering (NJ). The common ancestor of the entire IHF/HU family lies with HU sequences in both the PCA and the NJ plots (Figure 3B, C). As a control, the last common ancestors of the individual subunits lie within clusters representing that subunit. The common ancestor of all IHFs lies with IHFβ.

Thus, both ancestral state reconstruction of the presence/absence states of the various IHF/HU family subunits and ancestral sequence reconstruction on a TraM-rooted gene tree suggests that the TF IHF evolved from the non-specific DNA binding and nucleoid-shaping HU after the latter had diversified.

### Both IHF subunits have acquired residues for DNA binding in distinct ways

Now that we have characterised the nature of the overall common ancestor of the family, we asked how residues known to be involved in DNA binding have emerged along the lineage. We assembled a list of such residues from various structural and mutational studies (Goshima et al., 1990; Köhler & Marahiel, 1998; Liao et al., 2017; Lee et al., 1992; Mengeritsky et al., 1993; Pavlik & Spidlova, 2022; Read et al., 2000; Rice et al., 1996; Yoshua et al., 2021; Chen et al., 2004). Whereas data for HU were derived from studies on multiple organisms, that for IHF was limited to those from *E. coli* (Supplementary Table 1). This list of functional residues had been derived using a range of experimental techniques, including Atomic Force Microscopy, proteomics, site directed mutagenesis studies with challenge phage assays, EMSA, ChIP-seq analysis and DNAse 1 footprinting analysis and computational molecular dynamics simulations. As highlighted in a recent study concerning (Yoshua et al., 2021), IHF heterodimers can adopt 3 different DNA bending modes, with the “fully wrapped” form making base-specific contacts with DNA. The maximum DNA bend observed in the IHF-DNA complex (147°) is also a characteristic of the fully-wrapped form and hence, residues making base specific contacts in this conformation may also have a role to play in DNA bending.

We first tested the conservation of these experimentally determined residues of functional importance. To do this, we defined residues conserved in each subunit from our alignment using Shannon’s entropy (See Methods; Supplementary Table 1). Of 30 key residues from the literature, 18 are conserved in a varied distribution across the 4 subunits. While some residues show patterns of conservation specific to individual subunits (Lys86 of HU; Thr4, Ser47 of IHFα; Arg42, Glu44, Arg46, Val70 of IHFβ), most of the residues important for DNA binding are shared across different subunit classes. For instance, Arginine in position 126 of the alignment and Proline in position 136 are involved in DNA binding in multiple subunits and conserved universally (Arg58 in HU and Arg60 in IHFα; Pro64 in HU, Pro65 in IHFα, Pro64 in IHFβ respectively). Some residues known to facilitate DNA binding in only one subunit are, in fact, conserved across multiple subunit proteins. The Lys5 and Ile73 residues for IHFα are conserved in both IHFα and HU, while Phe47 and Arg61 from HU are conserved in all the subunits. We also noticed that Lys20 of IHFα is conserved in only HU, and Lys80 from HU is conserved in IHFβ.

To get an understanding of how and when these residues might have emerged, we calculated a Position-specific scoring matrix (PSSM) (See Methods) based on all the literature-derived residues and scored all sequences based on this matrix. HU residues known to bind to DNA are not conserved in early branching HU ancestral clades of the gene tree but are strongly conserved in late branching clades, with weak conservation in the ancestors of IHF subunits. Some of these residues appear to be conserved in IHFα but not in IHFβ. Key residues of IHFα (both in terms of general DNA binding as well as specific base contacts), however, show very strong conservation right from the early branching ancestral HU clades as well as the direct IHFα ancestors, with weak conservation in late branching HU and IHFβ clades. Residues known to bind to DNA in IHFβ are restricted only to the common ancestors of IHFβ and do not show conservation in any other clades. (Figure 4). IHFα-like proteins show a similar characteristic residue pattern as IHFα. However, the small number of proteins (N=34) in the IHFβ lineage in *Acidobacteria* appears to conserve few of the residues characteristic of the subunit as defined by the Proteobacterial lineage, but instead retains more HU-like properties (Supplementary Figure S9). PSSM scores calculated on the structure-guided alignment produced results consistent with those described above (Supplementary Figure S10).

**Figure 4.**
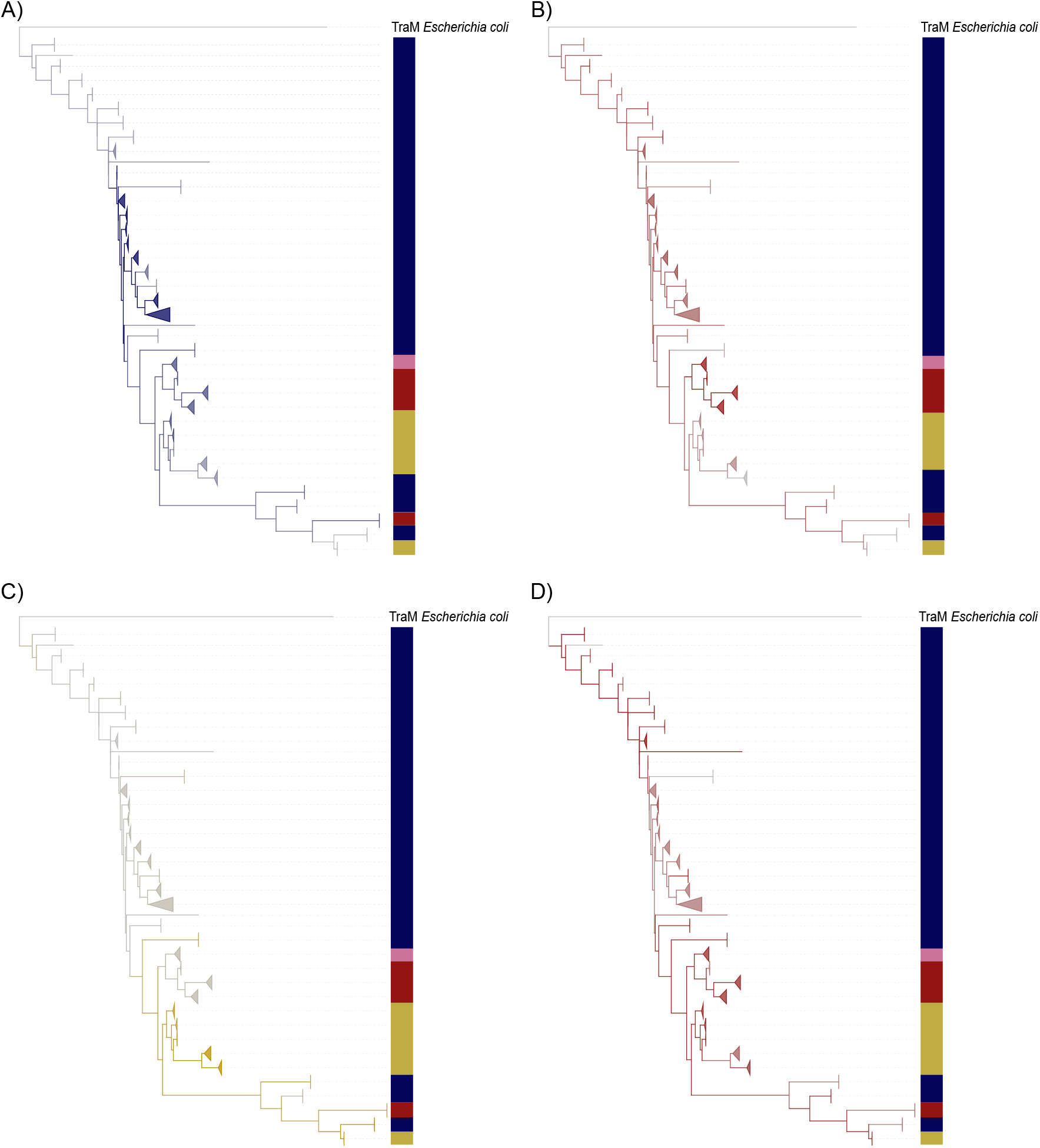
Conservation of Key residues. Collapsed IHF/HU rooted gene trees coloured on the basis of PSSM scores calculated for key residues important for DNA binding in (A) HU proteins (α and β), (B) IHFα, (C) IHFβ, and (D) Key residues important for making specific base contacts by IHFα in fully wrapped form (Yoshua et al., 2021). Colourstrips annotation indicates nature of sequences in collapsed gene tree (visualized using iTOL).

Thus, most DNA binding residues of IHFβ have emerged independently in the proteobacterial lineage but not in *Acidobacteria*. However, a proportion of DNA binding residues of IHFα have likely been co-opted from HU and gained the ability to bind to DNA in a sequence-specific manner only in IHF dimers. Whether IHFα-like in *Bacteroidetes* also forms sequence-specific DNA contacts in the absence of an IHFβ is to be tested. Thus, each of the IHF subunits have undergone different trajectories to acquire their DNA binding properties.

## Discussion and Conclusion

In summary, we present the following model for the evolution of the IHF/HU sequence family (Figure 5). HUβ, known to be a non-specific DNA binding protein involved in chromosome shaping, being the most conserved among the four subunits, was likely present in the last bacterial common ancestor, and is the only representative of this family in a large portion of the bacterial tree. That HUβ is ancestral is also supported by the IHF/HU gene tree rooted with TraM as the outgroup. IHFβ emerged from an HUβ-like sequence. Whether this was a duplication of HUβ in that ancestral node or an acquisition of a second copy of HUβ from some other source is unclear. This event probably occurred in the last common ancestor of *Proteobacteria* and *Acidobacteria*, which is the earliest node supporting consistent presence of IHFβ in downstream, extant nodes. The notion that IHFβ emerged before IHFα is also supported by the clustering of the common ancestor of all IHFs with IHFβ. This IHFβ appears to have diverged along distinct lines in the Proteobacterial and Acidobacterial lineages. The Acidobacterial IHFβ appears to retain more of the characteristic properties of HU than the Proteobacterial IHFβ. This might be explained purely by speciation, and/or by the fact that the Acidobacterial lineage does not contain an IHFα for its IHFβ to heterodimerise with, unlike in the Proteobacterial line. Whether this protein heterodimerises with HU, or homodimerises to bind DNA, and whether it does so in a sequence-specific manner, remains to be seen.

**Figure 5.**
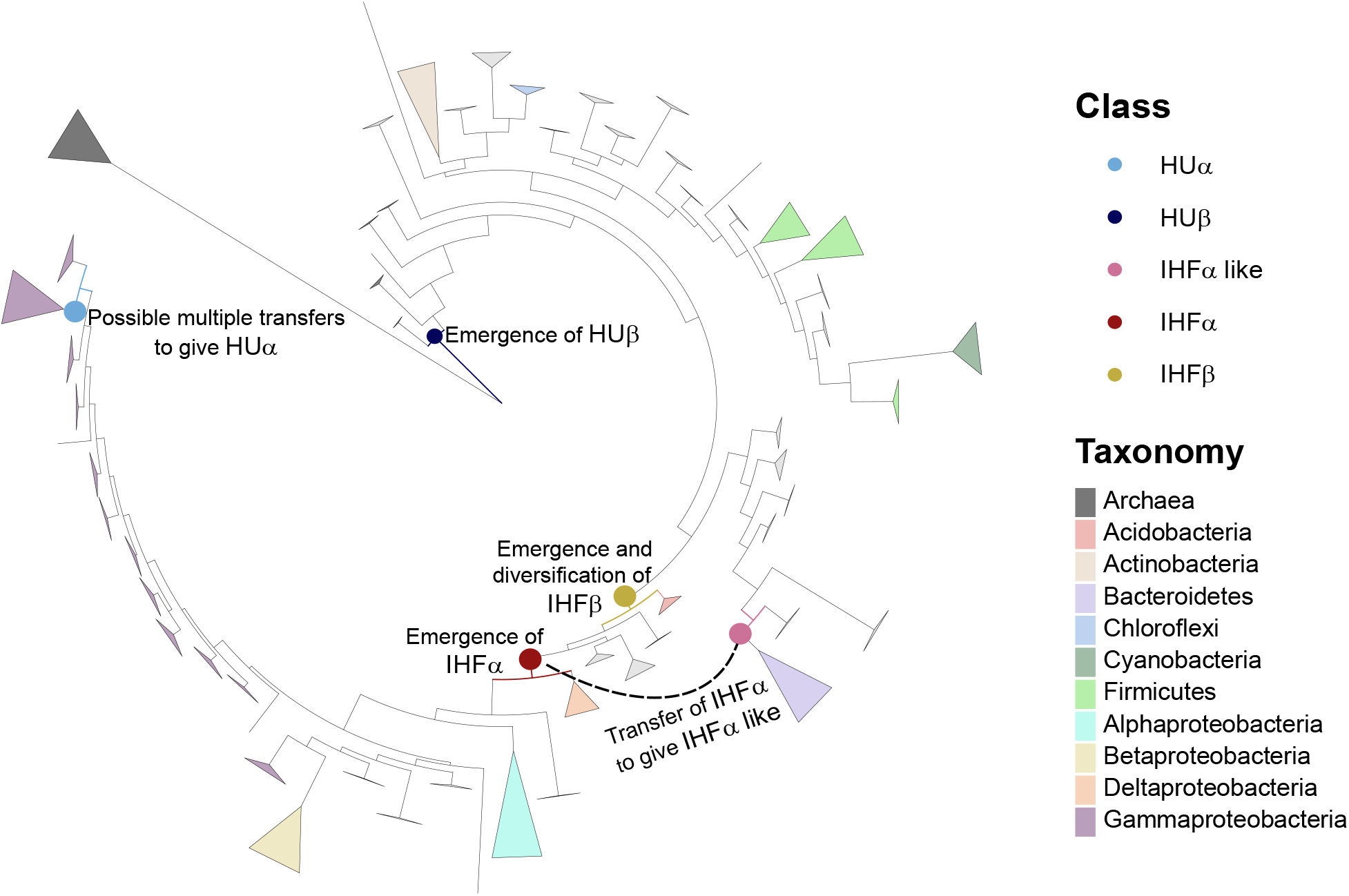
Evolution of the IHF/HU family. A model for the evolution of the IHF/HU sequence family on a bacterial species tree.

Soon after the formation of IHFβ, and before it gained its characteristic residue composition, another event along the lineage towards the *Proteobacteria*, possibly a duplication of IHFβ, produced IHFα. The two subunits of IHF must have co-evolved to dimerise and function as a cognate pair with DNA bending and TF activity. IHFα appears to have co-opted residues present in HU, plus other residues, to bind to DNA and to do so specifically in a manner that HU does not. That HU has some limited ability to recognize kinked DNA and act as a TF at a few sites may arise from the presence of such residues (Verma et al., 2023). In this context, we also highlight a recent study that showed that IHF heterodimer also has multiple DNA-binding ‘modes’ of which the so-called “fully-wrapped” conformation is the one in which the protein forms base-specific contacts through IHFα (Yoshua et al., 2021). IHFβ, on the other hand, has elaborated a distinct set of characteristic residues and lost most residues characteristic of HU. IHFα-like, which is present exclusively in *Bacteroidetes* and forms a sister-clade to IHFα within the larger IHF clade, might have emerged by horizontal transfer from an IHFα from the Proteobacterial lineage. This is supported by the inference that the node at which Proteobacterial IHFα is first gained is closer to the root but not ancestral to the node at which IHFα-like is first inferred to have appeared. Given that the *Bacteroidetes* do not code for IHFβ, this transfer event might have occurred before IHFα and IHFβ co-evolved to dimerise in *Proteobacteria*. Like the *Acidobacterial* IHFβ, it is unknown whether IHFα-like also homodimerizes or heterodimerizes with HUβ. One can also ask whether IHFα-like (and *Acidobacterial* IHFβ) show specificity intermediate between classical HU and IHF.

A recent computational analysis of the sequences of IHF/HU family proteins in *Bacterioidetes* has suggested that these proteins may be related to certain eukaryotic proteins involved in chromosome partitioning (Burroughs et al., 2017). A fraction of these proteins have more than one copy of the HU domain and acquired additional domains such as FtsN and LysM, which might tether the protein to the cell surface with the DNA binding domain located in the cytoplasm. According to Burroughs et al., these proteins might help tether the nucleoid to the cell surface in *Bacteroidetes*, but may be involved in protein-protein interactions in eukaryotes. These proteins are likely products of lineage-specific expansions of this family in *Bacteroidetes*. Most of these multidomain examples (defined by the presence of a second domain or by long stretches without a domain annotation) of Bacteroidetes proteins are not part of our dataset (N = 916). This is because these are not annotated in orthology databases as IHF or HU, a filter we had used in assembling our dataset. However, a proportion of the single domain IHFα-like proteins characteristic of *Bacteroidetes* in our dataset do retain the signature – defined by the presence of an overlapping sequence family called “HU-HIG” in the Pfam database – described by Burroughs and colleagues. And the IHF/HU domain present in *Bacteroidetes* proteins not included in our dataset (a) does not cluster with IHFα-like, (b) scores poorly on PSSMs for known key residues in HU and IHF, (c) does not form an outgroup to the HU/IHF family, suggesting that it form a distinct lineage within IHF/HU (Supplementary Figure S11).

HUα appeared in a Gammaproteobacterial ancestor, thus gaining the ability to form a variety of homodimers and heterodimers with distinct structural and functional properties (Claret & Rouviere-Yaniv, 1997; Hammel et al., 2016). The formation of HUα might have been through transfer events rather than duplication, given that HUα sequences do not necessarily cluster with Gammaproteobacterial HUβ but with HUβ from various other clades. The sporadic occurrence of IHFα or IHFβ in early-branching bacterial lineages might be a consequence of horizontal transfer or errors in the phylogeny (for example, *Nitrospirae*, which appears as an IHFα/IHFβ-containing early-branching lineage in the 16S rRNA tree, clusters within *Deltaproteobacteria* in the GTDB tree). Finally, similar to the expectation for a large majority of bacterial gene families (Treangen & Rocha, 2011), the IHF/HU family appears to have been subjected to extensive horizontal gene transfer events, as inferred from as many as 1415 pairs of within-subunit transfer events identified by gene tree-species tree reconciliation analysis (See Methods; N_speciation_ =1094, N_duplication_ = 124).

Thus, the sequence-specific DNA binding TF IHF likely evolved from a non-specific nucleoid-shaping protein HU. This, however, does not necessarily mean that this idea is a general principle applicable to a broad range of TF families. Additionally, the evolution of sequence specificity and the ability to recruit RNA polymerase and respond to a signal might also require the evolution of additional DNA binding or protein-protein / protein-ligand binding domains and interfaces. How these have shaped the evolution of transcriptional regulatory networks in prokaryotes and eukaryotes is an open question.

## Data Availability

All necessary data is available at http://doi.org/10.6084/m9.figshare.28524998.v2.

Krishnaswamy, Madhumitha (2025). IHF/HU phylogenetic analysis dataset. figshare. Dataset. https://doi.org/10.6084/m9.figshare.28524998.v2

Supplementary Figures and Table are also available to download.

## Acknowledgements

We thank Sunil Laxman, Dasaradhi Palakodeti, Varadharajan Sundaramoorthy, Inder Raj Singh, Ashitha Arun, Ganesh Muthu, and Akshara Dubey for discussions on the project.

