## Supplementary Figures and Table for "The evolution of sequence specificity in a DNA binding protein family"

### Supplementary Data

#### Supplementary Figures

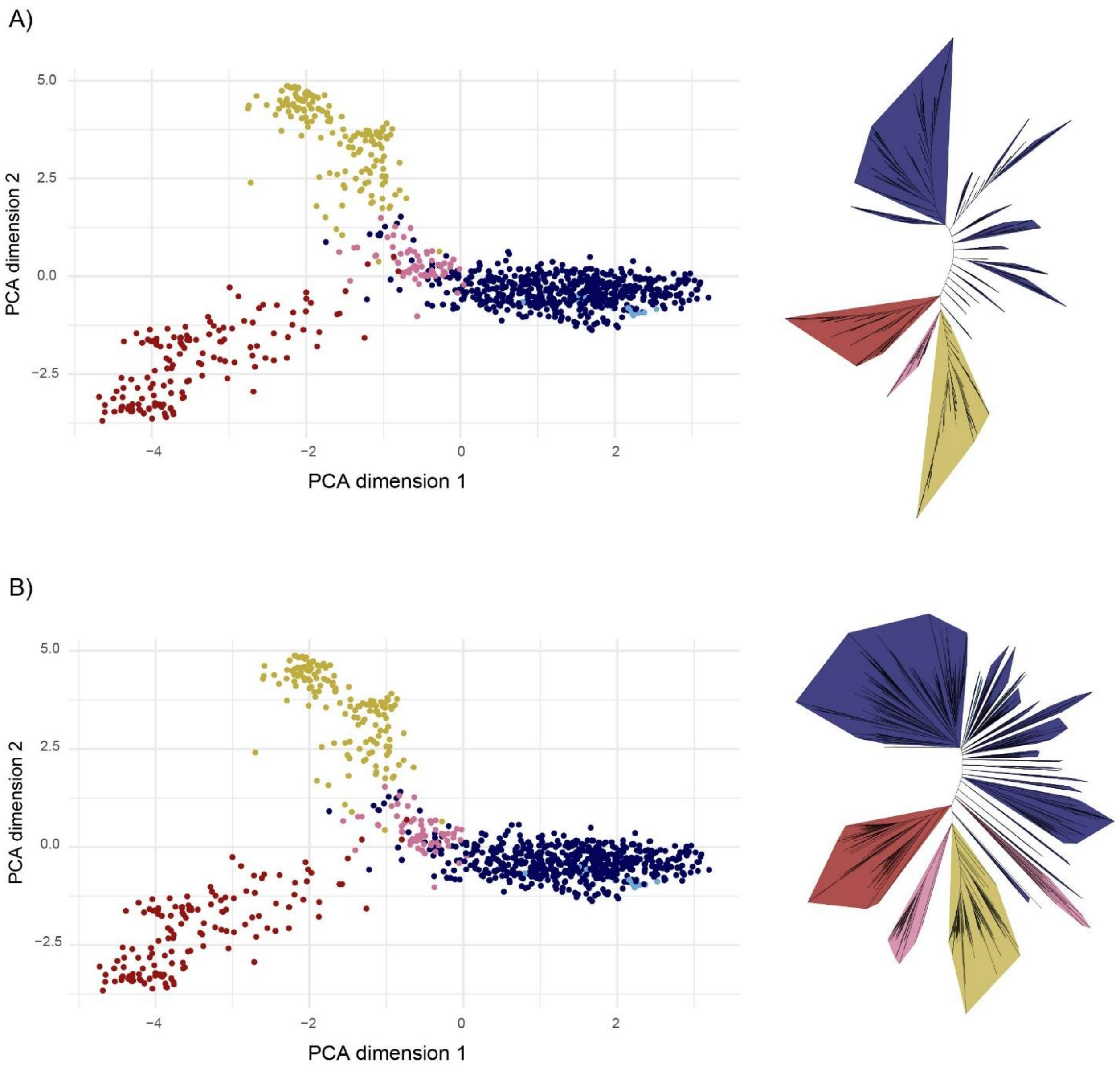


**S1:** PCA and NJ showing the distribution of (A) Subsampled sequences (N=990) taken from the original alignment (B) Subsampled sequences aligned guided by structure using MAFFT-DASH. Colouring scheme same as in main text and NJ trees visualized using iTOL.

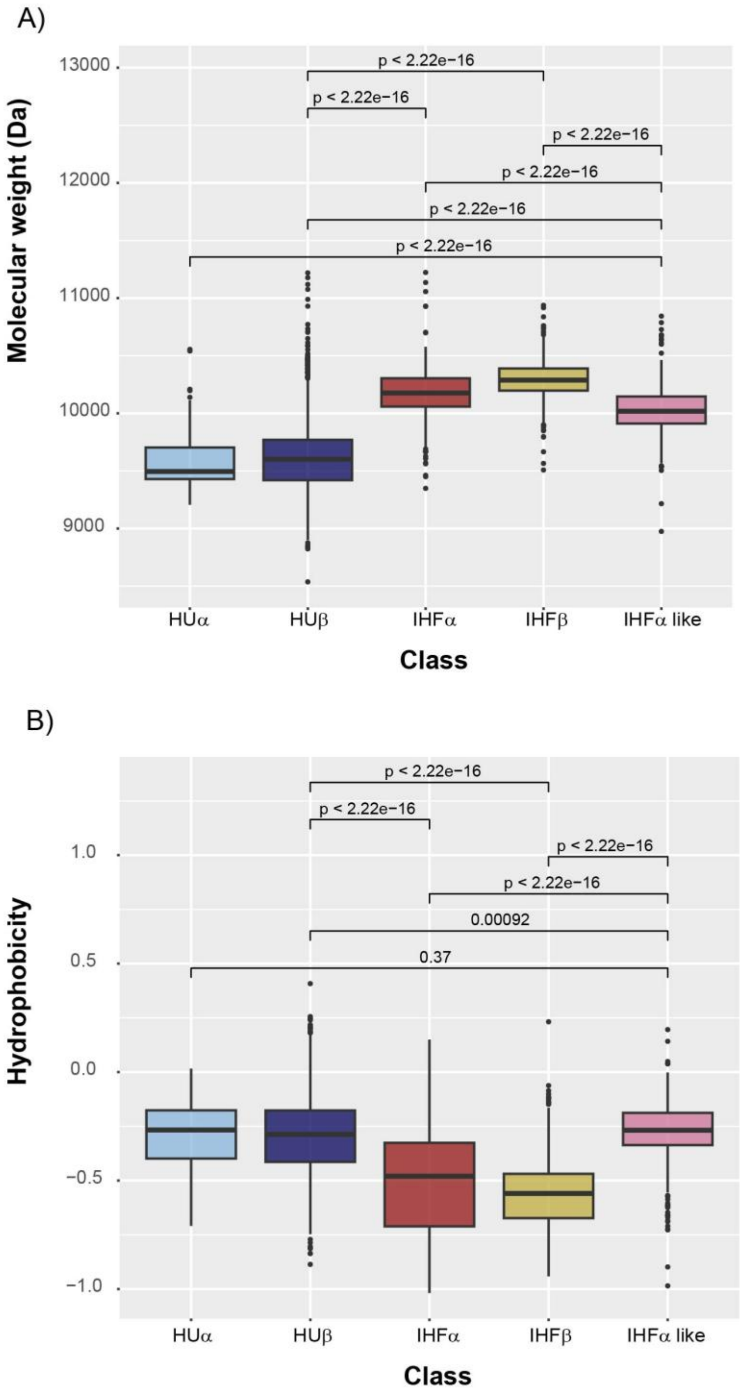


**S2:** Boxplot showing the variation of (A) Molecular weight and (B) Hydrophobicity between the different subunits. Molecular weights calculated using BioPython and Hydrophobicity calculated on Kyte and Doolittle hydrophobicity scale. Pairwise p values calculated using Wilcoxon test in R.

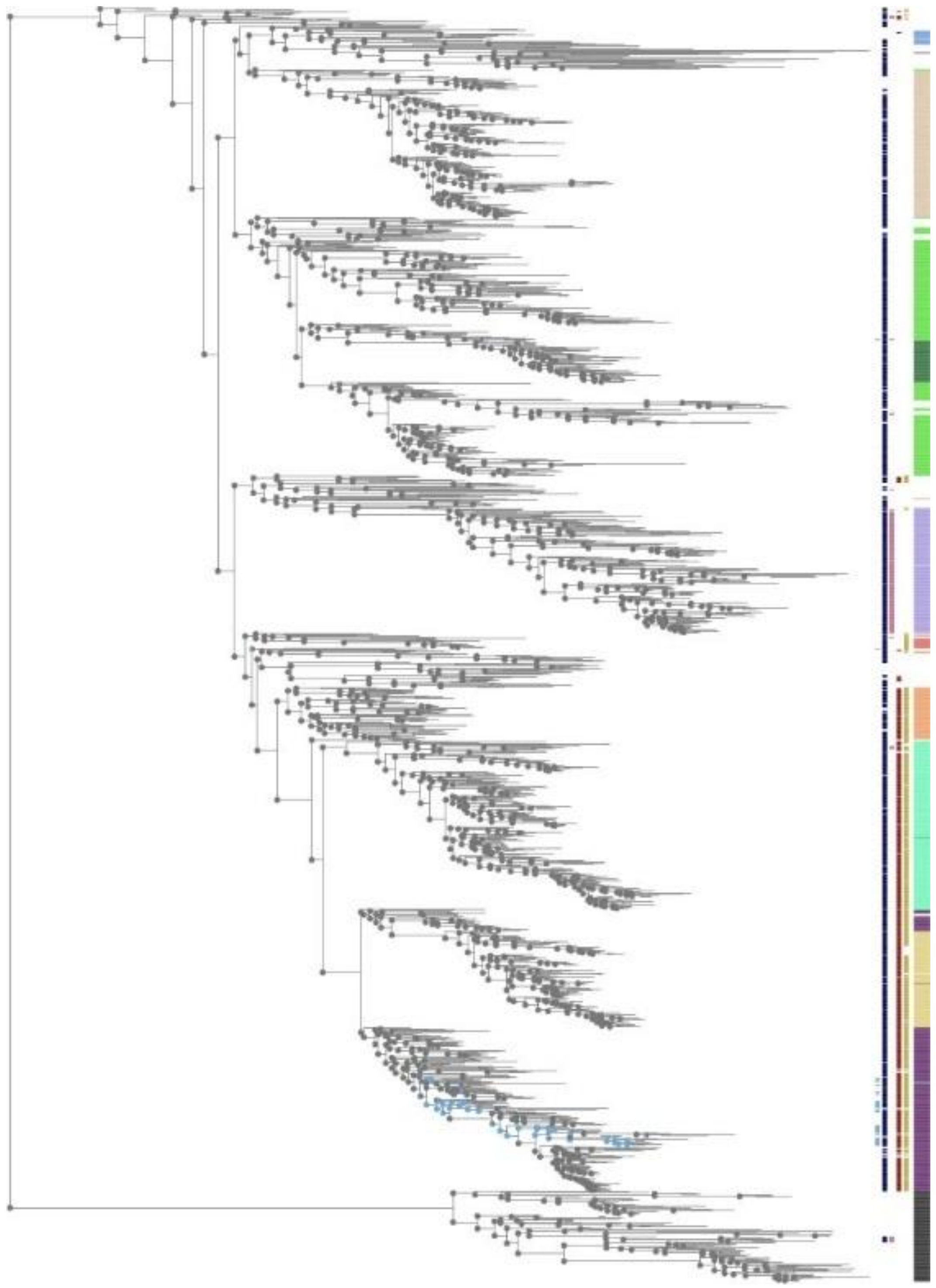

**S3:** Ancestral state reconstruction of HUα on the 16S rRNA tree. Colourstrips indicates extant presence absence, light blue circles indicate HUα presence. Visualized using iTOL.

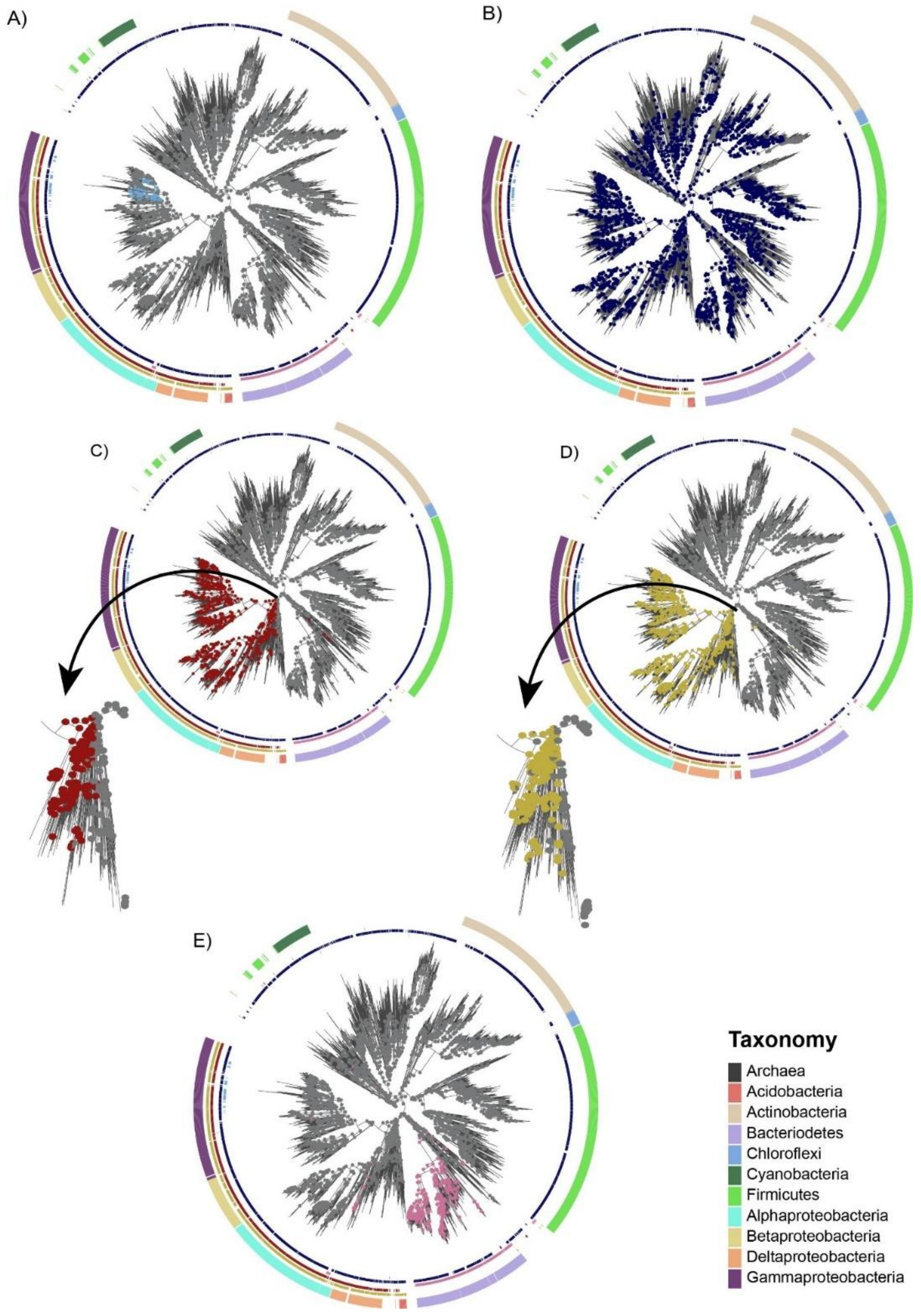


**S4:** Ancestral state reconstructions of all subunits on the GTDB tree with 3100 bacterial tips. (A) HUα (light blue), (B) HUβ (dark blue), (C) IHFα (red), (D) IHFβ (yellow), (E) IHFα-like (pink). Arrow marks for (B) and (C) emphasise the difference in first emergence for IHFα and IHFβ, consistent with 16S rRNA tree analysis.

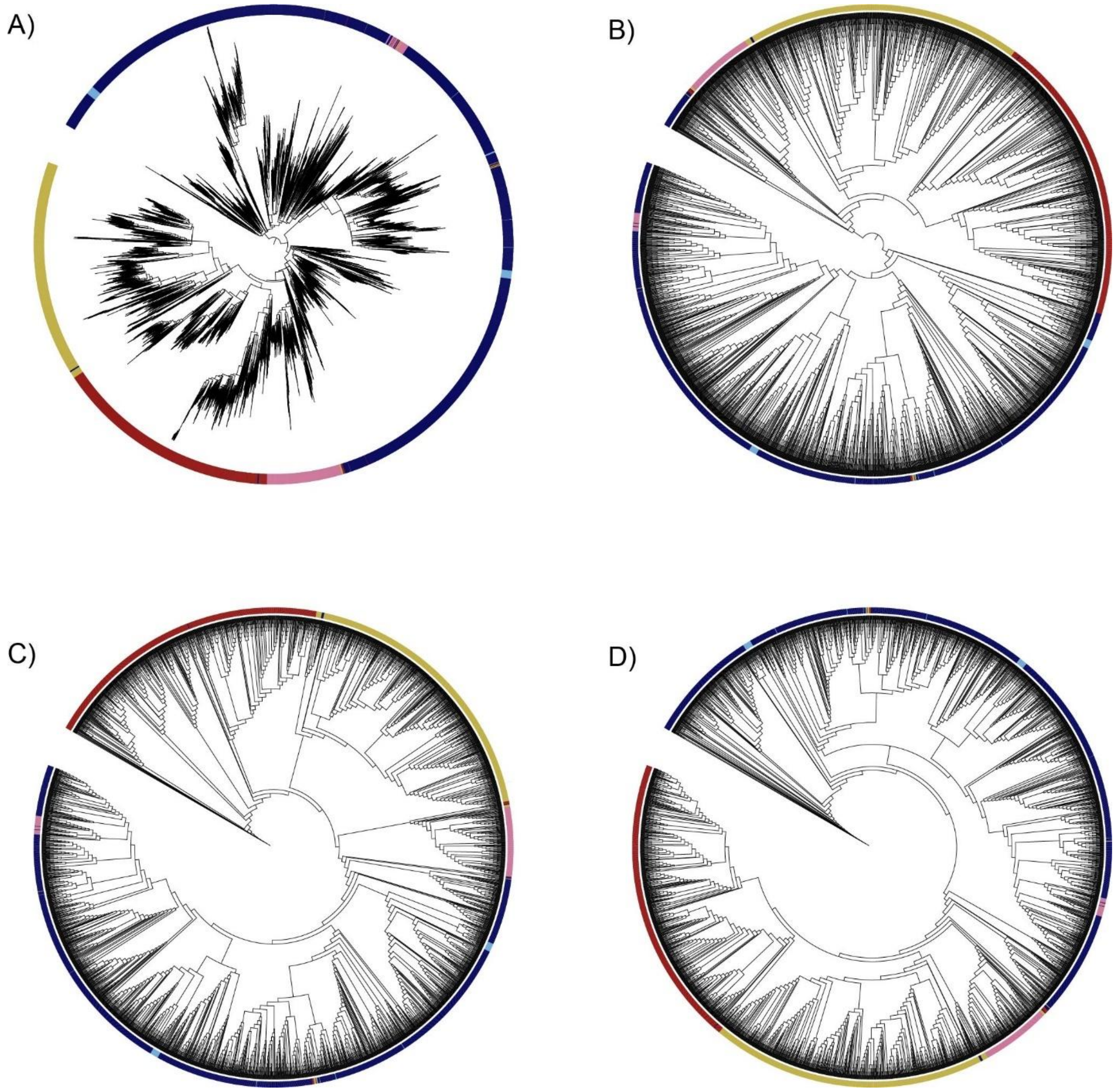


**S5:** Inconsistencies with midpoint rooting and OptRoot trees (A) Midpoint rooted tree (B) Subsampled midpoint rooted tree (C) OptRoot rooted tree (variable cost 1), (D) OptRoot rooted tree (variable cost 2)

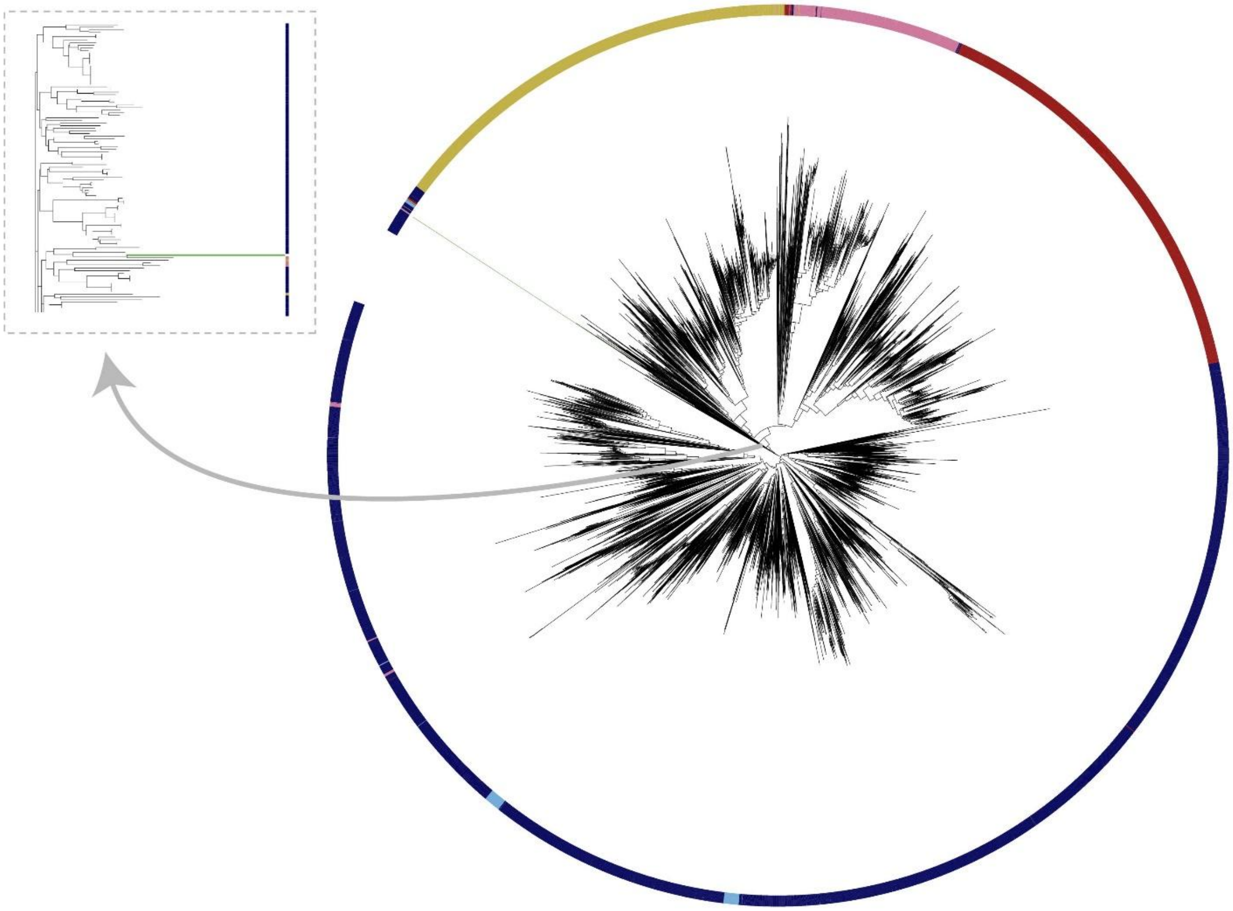

**S6:** Other outgroups - NJ of complete alignment with another member of PF00216 (PiliB) protein (outgroup highlighted in green). Inset zooms in on the tips close to the proposed outgroup (highlighted in green).

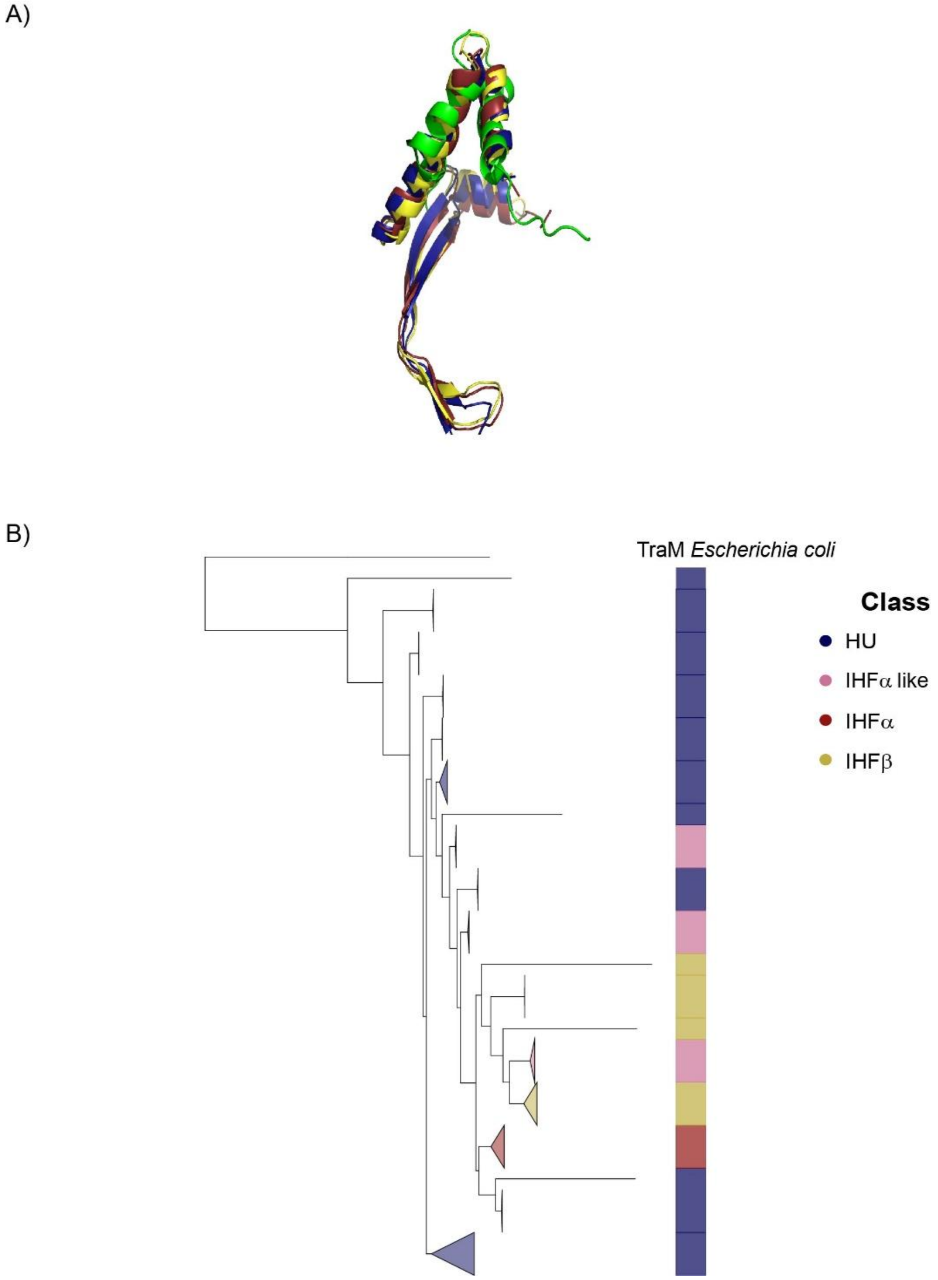


**S7:** (A) Structural superimposition of TraM (Green, PDB ID: 1DP3) with HU (Blue, PDB ID: 1P51), IHFα (Red) and IHFβ(Yellow) (PDB ID: 1IHF for both subunits). RMSD between TraM and HU: 0.873. Visualized using PyMol. (B) Neighbour joining (NJ) tree of TraM with the structure guided subsampled alignment, visualized using iTOL.

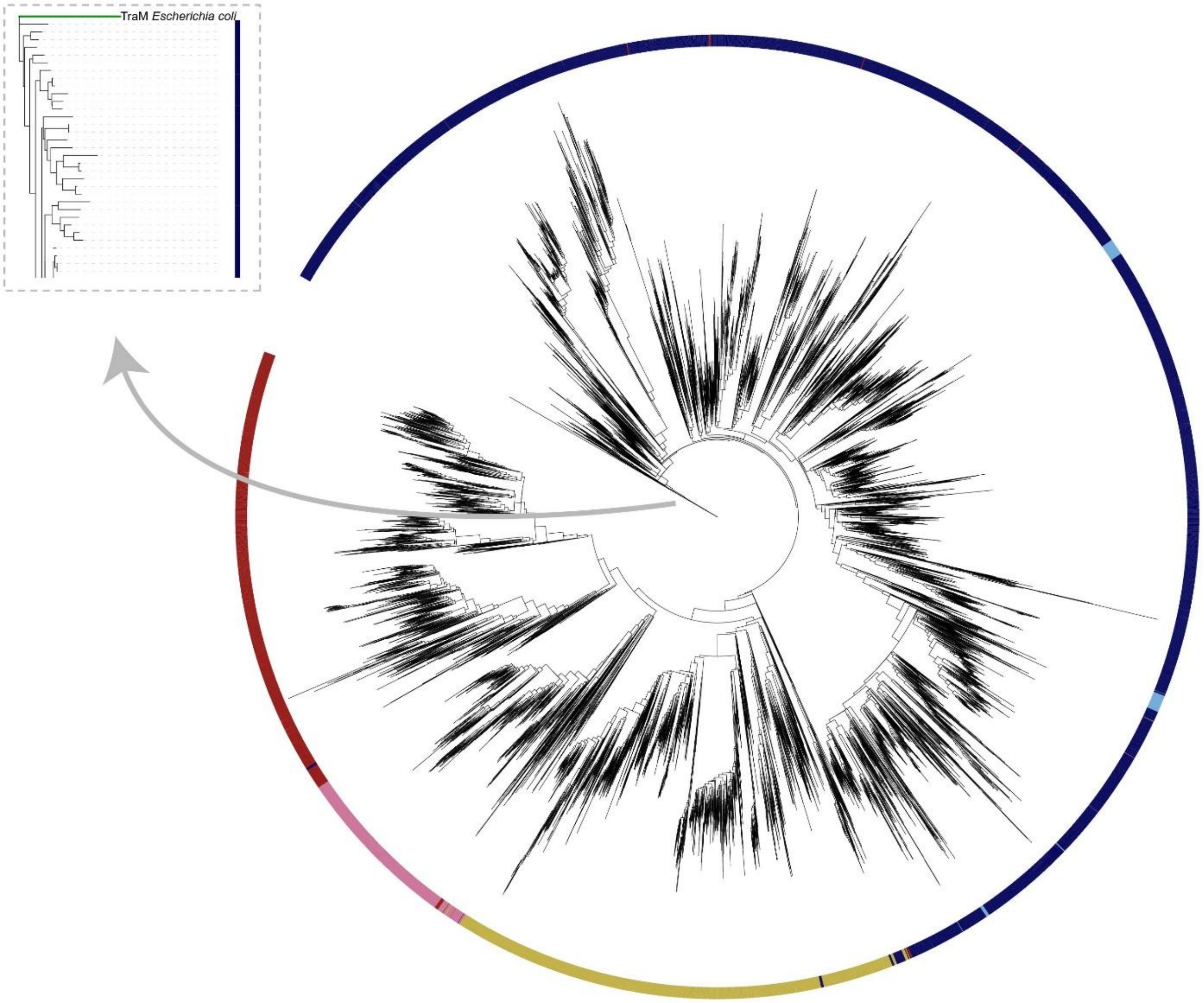


**S8:** Complete gene tree of all 9,880 IHF/HU domains rooted at TraM. HUα: Light blue, HUβ: Dark Blue, IHFα: Red, IHFβ: Yellow, IHFα-like: Pink. Visualized using iTOL. Inset highlights TraM (coloured in green) segregating from the extant sequences.

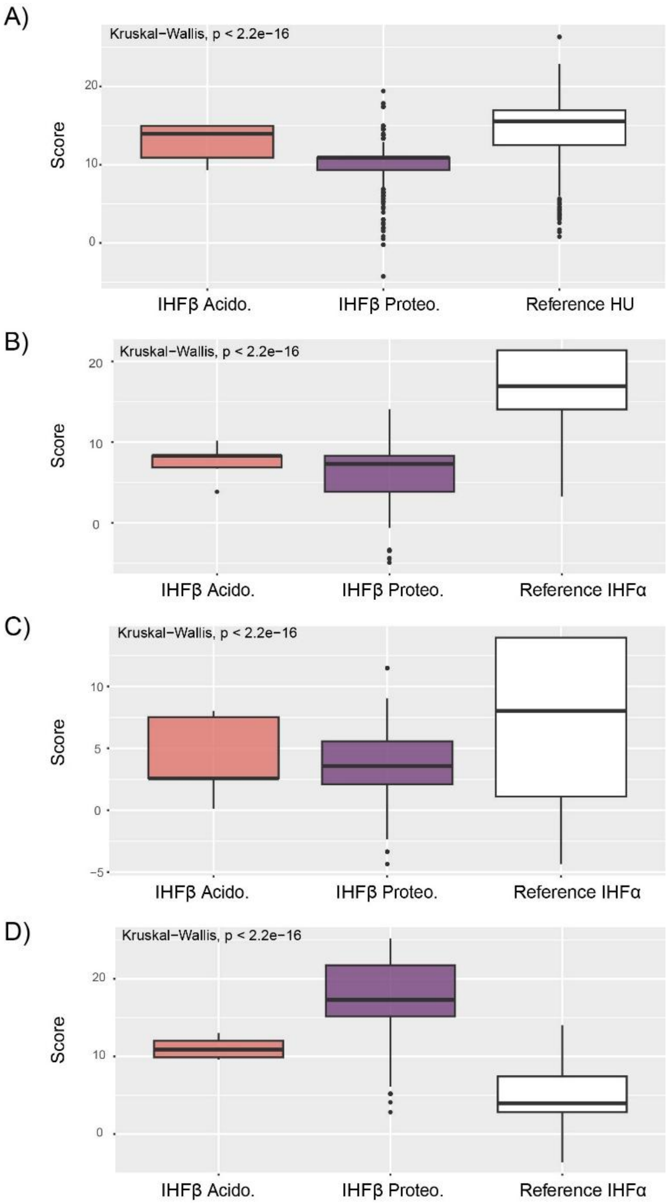


**S9:** Boxplot showing distribution of PSSM scores of IHFβ proteins between *Acidobacteria* (N=34) and *Proteobacteria* (N = 1432). (A) PSSM scores of HU key residues compared to reference HU in *Proteobacteria*, (B) PSSM scores of IHFα key residues compared to reference IHFα in *Proteobacteria*, (C) PSSM scores of specific base contact residues of IHFα compared to reference IHFα from *Proteobacteria*, (D) PSSM scores of IHFβ key residues.

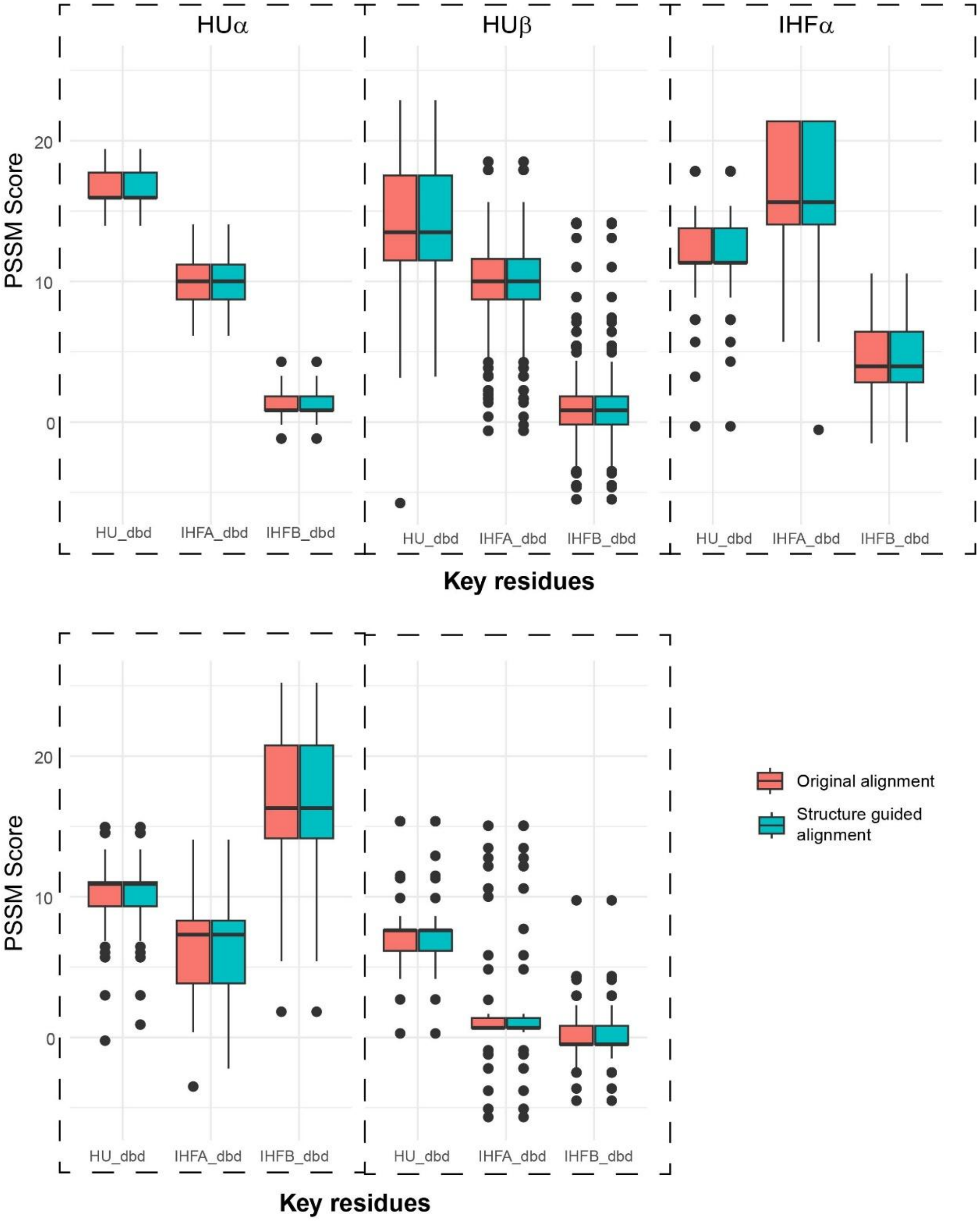


**S10:** Boxplots showing distribution of PSSM scores for key residues (x axis) compared for the subsampled original alignment (N=990) vs the structure guided alignment (N=990), with different plots for each of the different extant subunit classes

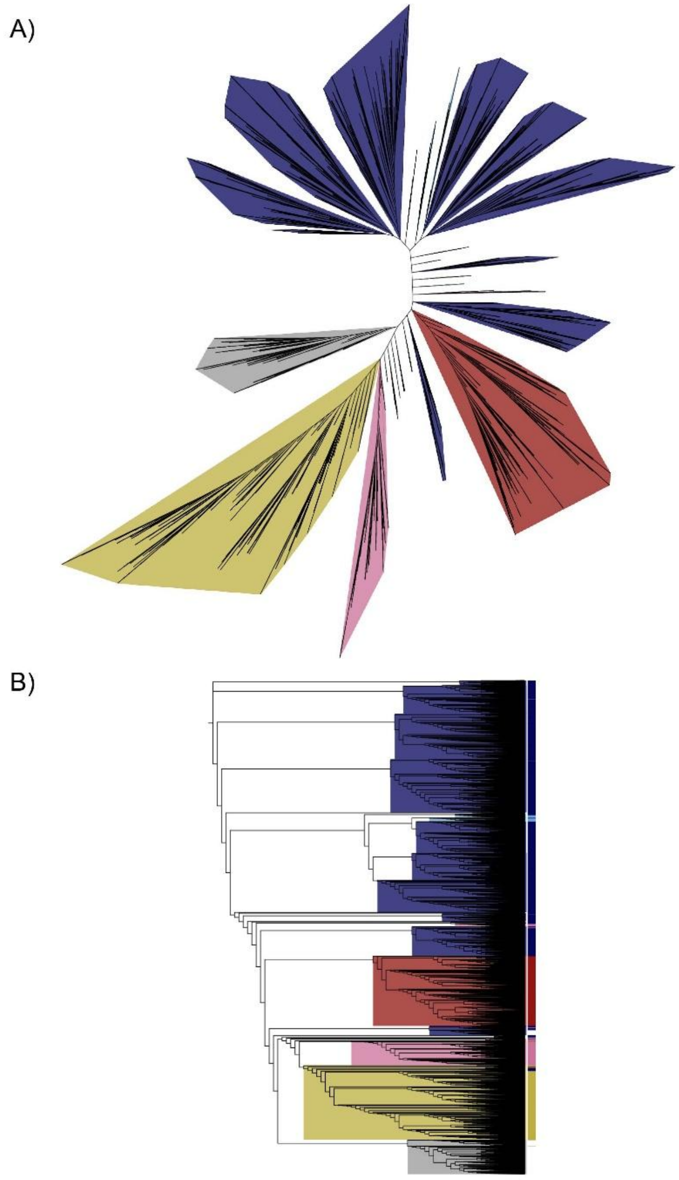


**S11:** Neighbour Joining (NJ) tree of the complete alignment (N=9880) (coloured as follows: HUα - light blue , HUβ - dark blue , IHFα - red, IHFβ - yellow, IHFα-like - pink) along with the domains of sequences (coloured grey) removed during curation (N=916) due to missing KO annotations, presence of multiple domains or extremely long sequence length.(A) Unrooted and (B) Rectangular tree showing that the grey clade forms a lineage within the IHF/HU family.

### Supplementary Table

| Full Alignment Position | Major residue in Alignment | Subunit | Residue | Reference species | Conservation in different subunits (conservation cutoffs) | | |
|---|---|---|---|---|---|---|---|
| | | | | | HU (0.359) | IHFα (0.2425) | IHF β (0.194) |
| 2 | Asn (47.49%) | IHFα | Thr4* | *Escherichia coli* [a] | No | Yes (0.084) | No |
| 3 | Lys (87.49%) | IHFα | Lys5* | *Escherichia coli* [a] | Yes (0.1510) | Yes (0.2424) | Yes (0.1889) |
| 28 | Lys (37.49%) | HU | Lys13 | *Acinetobacter baumannii* [b] | No | No | No |
| 52 | Lys (68.42%) | IHFα | Lys20* | *Escherichia coli* [a] | Yes (0.3047) | No | No |
| 53 | Lys (39.24%) | IHFα | Arg21* | *Escherichia coli* [a] | No | No | No |
| 56 | Glu (29.46%) | IHFα | Lys24* | *Escherichia coli* [a] | No | No | No |
| 95 | Lys (30.66%) | IHF β | Arg42 | *Escherichia coli* [c] | No | No | Yes (0.1705) |
| 100 | Glu (16.39%) | IHF β | Glu44 | *Escherichia coli* [c,d,e] | No | No | Yes (0.0211) |
| 103 | Val | IHFα | Ser47 | *Escherichia coli* [c,d] | No | Yes | No |

| | | | | | | | |
|---|---|---|---|---|---|---|---|
| | (24.35%) | | | | | (0.1413) | |
| | | IHF β | Arg46 | *Escherichia coli* [c] | No | No | Yes (0.0131) |
| 108 | Phe (90.14%) | HU | Phe47 | *Escherichia coli* [f] | Yes (0.2298) | Yes (0.0067) | Yes (0.0637) |
| 126 | Arg (86.54%) | HU | Arg58 | *Escherichia coli* [f] ; *Bacillus subtilis* [g] ; *Francisella tularensis* [h] | Yes (0.1556) | Yes (0.0057) | No |
| | | IHFα | Arg60 | *Escherichia coli* [c,d] | | | |
| 128 | Thr (28.26%) | IHFα | Pro61 | *Escherichia coli* [c,d] | No | No | No |
| 130 | Arg (84.83%) | HU | Arg61 | *Escherichia coli* [f]; *Bacillus subtilis* [g] ; *Francisella tularensis* [h] | No** (0.3688) | Yes (0.0322) | Yes (0.1259) |
| 135 | Asn (96.03%) | HU | Val63 | *Helicobacter pylori* [i] | No | No | No |
| 136 | Pro (93.68%) | HU | Pro64 | *Helicobacter pylori* [i] | Yes (0) | Yes (0.0281) | Yes (0.0259) |
| | | IHFα | Pro65 | *Escherichia coli* [c,d] | | | |
| | | IHF β | Pro64 | *Escherichia coli* [c] | | | |
| 138 | Lys | IHFα | Lys66 | *Escherichia coli* [c,d] | No | No | No |

| | | | | | | | |
|---|---|---|---|---|---|---|---|
| | (31.19%) | | | | | | |
| 148 | Ile (58.92%) | IHFα | Ile71 | *Escherichia coli* [c,d] | No | No | No |
| | | IHF β | Val70 | *Escherichia coli* [c] | No | No | Yes (0.1755) |
| 155 | Ile (73.46%) | IHFα | Ile73 | *Escherichia coli* [c,d] | Yes (0.2288) | Yes (0.1905) | No |
| | | IHF β | Leu72 | *Escherichia coli* [c] | No | No | No |
| 161 | Lys (26.04%) | HU | Ser74 | *Francisella tularensis* [h] | No | No | No |
| 172 | Lys (58.17%) | HU | Lys80 | *Bacillus subtilis* [g] | No | No | Yes (0.1246) |
| 180 | Lys (69.16%) | HU | Lys86 | *Bacillus subtilis* [g] | Yes (0.2821) | No | No |
| | | IHF β | Arg87 | *Escherichia coli* [j] | No | No | No |
| 182 | Ala (39.40%) | IHF β | Arg89 | *Escherichia coli* [j] | No | No | No |
| 183 | Val (67.41%) | IHF β | Ala90 | *Escherichia coli* [j] | No | No | No |

* Are involved in binding to specific DNA base contacts – IHF was shown to bind to the DNA in multiple modes of which the so-called ‘fully-wrapped’ mode (Yoshua et al., 2021), and considered to represent specific DNA binding

** Borderline conservation

[a] (Yoshua et al., 2021) ; [b] (Liao et al., 2017) ; [c] (Rice et al., 1996) ; [d] (Lee et al., 1992) ; [e] (Read et al., 2000) ; [f] (Goshima et al., 1990) ; [g] (Köhler & Marahiel, 1998) ; [h] (Pavlik & Spidlova, 2022) ; [i] (Chen et al., 2004); [j] (Mengeritsky et al., 1993)

**Supplementary Table 1:** Residues known to bind to DNA in different IHF/HU subunits, and their conservation within each subunit clade. The alignment positions is in reference to the alignment file with the ancestral nodes and TraM outgroup.
